# Receptor-binding domain recombinant protein on alum-CpG induces broad protection against SARS-CoV-2 variants of concern

**DOI:** 10.1101/2021.07.06.451353

**Authors:** Jeroen Pollet, Ulrich Strych, Wen-Hsiang Chen, Leroy Versteeg, Brian Keegan, Bin Zhan, Junfei Wei, Zhuyun Liu, Jungsoon Lee, Rahki Kundu, Rakesh Adhikari, Cristina Poveda, Maria Jose Villar, Syamala Rani Thimmiraju, Brianna Lopez, Portia M. Gillespie, Shannon Ronca, Jason T. Kimata, Martin Reers, Vikram Paradkar, Peter J. Hotez, Maria Elena Bottazzi

## Abstract

We conducted preclinical studies in mice using a yeast-produced SARS-CoV-2 RBD subunit vaccine candidate formulated with aluminum hydroxide (alum) and CpG deoxynucleotides. This formulation is equivalent to the Corbevax™ vaccine that recently received emergency use authorization by the Drugs Controller General of India. We compared the immune response of mice vaccinated with RBD/alum to mice vaccinated with RBD/alum+CpG. We also evaluated mice immunized with RBD/alum+CpG and boosted with RBD/alum. Mice were immunized twice intramuscularly at a 21-day interval. Compared to two doses of the /alum formulation, the RBD/alum+CpG vaccine induced a stronger and more balanced Th1/Th2 cellular immune response, with high levels of neutralizing antibodies against the original Wuhan isolate of SARS-CoV-2 as well as the B.1.1.7 (Alpha), B. 1.351 (Beta), B. 1.617.2 and (Delta) variants. Neutralizing antibody titers against the B.1.1.529 (BA.1, Omicron) variant exceeded those in human convalescent plasma after Wuhan infection but were lower than against the other variants. Interestingly, the second dose did not benefit from the addition of CpG, possibly allowing dose-sparing of the adjuvant in the future. The data reported here reinforces that the RBD/alum+CpG vaccine formulation is suitable for inducing broadly neutralizing antibodies against SARS-CoV-2 including variants of concern.

## Introduction

More than two years into the COVID-19 pandemic, the world has now recorded over 6 million deaths and more than 450 million infections [1]. With unprecedented efforts from governments, non-profits, the academic and the industrial sector, vaccines have become widely available in the Northern hemisphere, however, many low- and middle-income countries (LMICs) in Africa, Asia, and South America still lack access to COVID-19 vaccines [2]. For example, the vaccination rate in the United States exceeds that of Africa’s two most populous countries, Nigeria and Ethiopia by 23 and 45-fold, respectively [3]. Consequently, we must realize that overall, only a small fraction of the people in the world have yet been fully vaccinated [4]. In addition, unequal access to vaccines and continued circulation of the virus has resulted in more and more SARS-CoV-2 variants emerging globally [5]. Some of these variants have raised concerns for their ability to at least partially evade the immune response acquired after natural infection with SARS-CoV-2 or even after vaccination with the first-generation COVID-19 vaccines [6]. There is an urgent need to bridge these gaps in vaccine equity and global distribution.

While mRNA, viral-vector vaccines, and whole-inactivated virus vaccines have been crucial tools in the first round of controlling COVID-19, they have limitations concerning cost, production capacity, as well as distribution and storage. Recombinant protein-based vaccines are therefore expected to play an important role in providing expanded coverage to populations that have yet to receive their immunizations [7]. Their advantages include ease of production, absence of freezer-chain requirements, and decades-long track records of safety with durable protection. For that reason, we and other groups have focused on recombinant protein platforms for our portfolio of neglected disease ‘antipoverty’ vaccines to prevent emerging and neglected infection such as schistosomiasis, hookworm, or Chagas disease affecting LMICs [8, 9].

The receptor-binding domain (RBD) of the viral spike protein is one of the main targets for COVID-19 vaccines, and our studies found that RBD-based subunit vaccines represent an effective strategy for the development of vaccines against SARS [10] and MERS [11]. With respect to COVID-19, for instance, an RBD-ferritin-nanoparticle vaccine (RFN), expressed in mammalian cell lines and adjuvanted with Army Liposomal Formulation QS-21 (ALFQ) elicited SARS-CoV-2 neutralizing antibodies and a Th1-biased T-cell response in various animal models. Moreover, cross-neutralization with this vaccine candidate stayed robust against the alpha and beta variants [12–14]. A similar construct generated through the SpyCatcher/SpyTag system elicited high titers of broadly neutralizing antibodies in mice [15, 16]. Likewise, an aluminum hydroxide-adjuvanted RBD-dimer (ZF2001) recombinantly produced in Chinese hamster ovary cell lines is currently in Phase 3 clinical trials and was shown to be well-tolerated and able to produce neutralizing antibody titers exceeding the levels found in convalescent serum, albeit, ideally, in a three-dose regimen [13].

We previously demonstrated that yeast-expressed RBD proteins (RBD219-N1C1, RBD203-N1), formulated with aluminum hydroxide (alum), stimulate virus-neutralizing antibodies, and induce primarily a Th2 cellular immune response in mice [17, 18]. As the next step, we optimized the alum-based formulation by reducing the protein doses as well as by incorporating in the formulation, at point of injection, a synthetic oligodeoxynucleotide [19] that mimics the effect of CpG motifs in bacterial DNA, stimulating TLR9 receptors to induce a stronger and more balanced Th1/Th2 response [20].

In humans, the Class B CpG1018 has a strong track record in the clinic and is part of the formulation of HEPLISAV-B, a licensed hepatitis B vaccine. Related to COVID-19, CpG1018 has been used to induce robust protection in a non-human primate study as part of an RBD-nanoparticle vaccine [21], and several CpG1018-adjuvanted protein vaccines have now entered the clinic [22]. As an equivalent to CpG1018, the Class B CpG1826 [23], has been shown in mice to strongly activate B cells and elicit TLR9-dependent NF-κB signaling, and only weakly stimulate IFN-α secretion [24].

Here, we report the result of preclinical studies in mice using the yeast-produced SARS-CoV-2 RBD subunit vaccine candidate formulated with aluminum hydroxide (alum) and CpG1826 deoxynucleotides. We compared the humoral responses, the vaccine-induced neutralizing antibody titers, and the cellular immune responses of mice vaccinated with RBD/alum to that of mice vaccinated with RBD/alum+CpG. In addition, we also evaluated the option to prime the mice with RBD/alum+CpG and boost with RBD/alum alone. This formulation is equivalent to the Corbevax™ vaccine that recently received emergency use authorization by the Drugs Controller General of India [25].

## Materials and Methods

### Sources of recombinant protein RBD219-N1C1 and RBD203-N1

The recombinant SARS-CoV-2 RBD219-N1C1 protein was designed and produced originally at Baylor College of Medicine, using previously published methods [26, 27]. Lot# S2RBD110620JXL-1 (formulated with alum) was used as a reference protein. Using the same master cell bank, the recombinant protein was produced at scale by Biological E (Hyderabad, India) and shipped to Baylor College of Medicine (BCM) (lot#BC07/001/20) where it was used in preclinical studies. A second version of the recombinant RBD protein, SARS-CoV-2 RBD203N1 was also produced at Baylor College of Medicine. SARS-CoV-2 RBD203-N1 is a truncated version of RBD219-N1C1, which after fermentation and purification had identical biophysical properties as RBD219-N1C1. Furthermore, in a side-by-side pre-clinical immunogenicity study, no statistical differences were observed between the two recombinant RBD proteins, and thus we consider them as equal [18]. SARS-CoV-2 RBD203-N1 was used in the second set of preclinical studies to test the neutralizing antibodies in sera of vaccinated mice against the Wuhan (D614G), Delta, and Omicron variants of the SARS-CoV-2 pseudovirus.

### Vaccine formulations

SARS-CoV-2 RBD proteins were diluted in 20 mM Tris, 100 mM NaCl, pH 7.5 (TBS buffer) before mixing with alum (Alhydrogel^®^, aluminum hydroxide, Catalog # 250-843261 EP, Croda Inc., UK). CpG1826 (Invivogen, USA) was added at point of injection.

### Preclinical Design

All animal experiments were performed under a Baylor College of Medicine Institutional Animal Care and Use Committee approved protocol. Female BALB/c mice, age 11-13 weeks old, were immunized twice intramuscularly at 21-day intervals with formulations shown in Table 1, and then euthanized two (2) weeks after the second vaccinations.

**Table 1:**
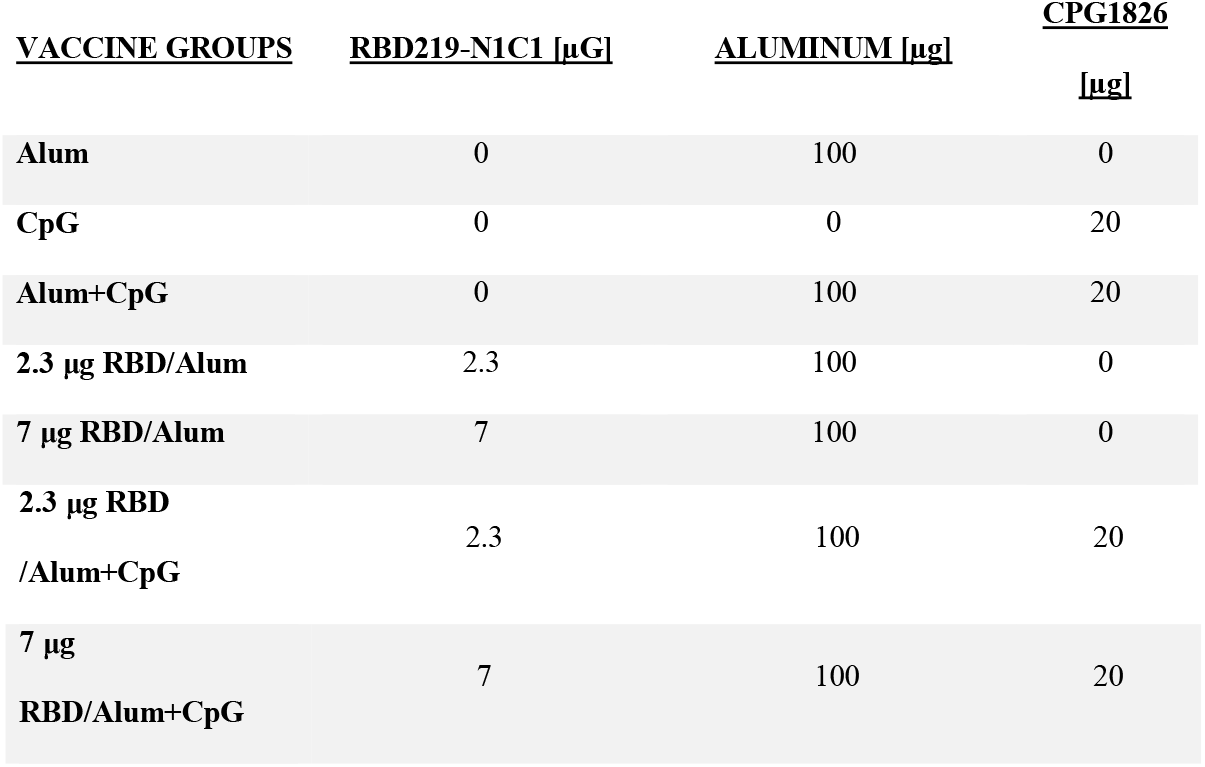
Vaccine formulation table

### Serological Antibody Measurements by ELISA

To examine SARS-CoV-2 RBD-specific antibodies in the mouse sera, indirect ELISAs were conducted as published elsewhere [17]. Briefly, plates were coated with 0.2 μg/well SARS-CoV-2 RBD219-N1C1. Mouse serum samples were 3-fold diluted from 1:200 to 1: 11,809,800 in 0.1% BSA in assay buffer (0.05% Tween-20 in 1x PBS). Assay controls included a 1:200 dilution of pooled naive mouse sera and assay buffer as blanks. 100 μL/well of either 1:6000 goat anti-mouse IgG HRP, 1:6000 goat anti-mouse IgG1 HRP, or 1:6000 goat anti-mouse IgG2a HRP in assay buffer was added. After incubation plates were washed five times followed by the addition of 100 μL/well 4°C TMB substrate. Plates were incubated for 15 minutes at room temperature (RT) while protected from light. After incubation, the reaction was stopped by adding 100 μL/well 1 M HCl. The absorbance at a wavelength of 450 nm was measured using a Biotek Epoch 2 spectrophotometer

The titer cutoff value was calculated using the following formula: Titer cutoff = average of negative control + 3 * standard deviation of the negative control. Then, for each sample, the titer was determined as the lowest dilution of each mouse sample with an average O.D. value above the titer cutoff. When a serum sample did not show any signal at all and a titer could not be calculated, an arbitrary baseline titer value of 67 was assigned to the serum sample. All data were plotted in Graphpad Prism 6 as individual samples and the geometric mean.

### Cytokine secretion in re-stimulated splenocytes by Luminex

A single-cell suspension from mouse splenocytes was prepared according to methods published elsewhere [17]. Briefly, spleens were dissociated using GentleMACS Octo Dissociator. Pelleted splenocytes were resuspended in 1 mL ACK lysing buffer by pipetting to lyse red blood cells. After 1 minute at RT, the reaction was stopped by the addition of 40 mL PBS. Tubes were centrifuged for 5 minutes at 300 x g at RT. Then supernatant was discarded and splenocytes were resuspended in 5 mL 4°C cRPMI (RPMI 1640 + 10% HI FBS + 1x pen/strep). Splenocyte suspensions were transferred through a 40 μm filter to a new 50-mL conical tube to obtain a single-cell suspension. For the *in vitro* cytokine release assay, splenocytes were seeded in a 96-well culture plate at 1×10^6^ live cells in 250 μL cRPMI. Splenocytes were (re-)stimulated with 10 μg/mL RBD219-N1-C1 protein, PMA/Ionomycin (positive control), or media (negative control) for 48 hours at 37° C 5% CO2. After incubation, 96-well plates were centrifuged to pellet the splenocytes down and supernatant was transferred to a new 96-well plate. A Milliplex Mouse Th17 Luminex kit (EMD Millipore) with analytes IL-1β, IL-2, IL-4, IL-6, IL-10, IL-13, IL-17A, IFN-γ, and TNF-α was used to quantify the cytokines secreted in the supernatant by the re-stimulated splenocytes according to a previously published method [28]. The readout was performed using a MagPix Luminex instrument. Raw data was analyzed using Bio-Plex Manager software, and further analysis was done with Excel and Prism. For the visualization of the individual cytokine results in the heatmap, the median was calculated of the cytokine concentrations of each group.

### Pseudovirus assay for determination of neutralizing antibodies

The pseudovirus used was a non-replicating lentiviral particle that has the SARS-CoV-2 spike protein on its membrane and can express a luciferase gene after infection. Using *in vitro* grown human 293T-hACE2 cells, infection was quantified based on the expression of luciferase [17].

The plasmids used for the pseudovirus production were the luciferase-encoding reporter plasmid (pNL4-3.lucR-E-, [29]), Gag/Pol-encoding packaging construct (pΔ8.9, [30]), and codon-optimized SARS-CoV-2 spike variants of concern protein expressing plasmids (pcDNA3.1-CoV-2 S gene) based on clone p278-1 [31]. Pseudovirions were generated by transfection of 293T cells as previously described [17]. Pseudovirus containing supernatants were recovered after 48 hours and passed through a 0.45 μm filter and saved at −80°C until used for neutralization studies.

In order to test the different pseudovirus variants, the following RBD mutations were introduced into a spike expression clone that includes the D614G mutation: N501Y (mimics the B.1.1.7 strain), and K417N, E484K, N501Y (mimics the B.1.351 strain). Mutations were generated sequentially by overlap extension PCR in 20 μL reactions using Phusion Green DNA polymerase master mix (ThermoFisher) according to the manufacturer’s protocol, 2 ng of the spike plasmid clone as template, and 500 nM of each PCR primer of a set in Table 2. The basic PCR cycling protocol was denaturation at 98°C for 30 sec followed by 25 cycles of 98°C for 10 sec, 30 sec at the annealing temperature, and extension at 72°C for 4 min 40 sec. A final elongation step at 72°C for 6 min was included. The annealing temperature for each primer set was determined using the ThermoFisher web-based Tm Calculator for Phusion DNA polymerase. PCR products were digested with DpnI for 1 hour at 37°C and NEB 10 beta cells were transformed and grown for plasmid isolation. The mutations in the spike coding region were confirmed by Sanger sequencing.

**Table 2.**
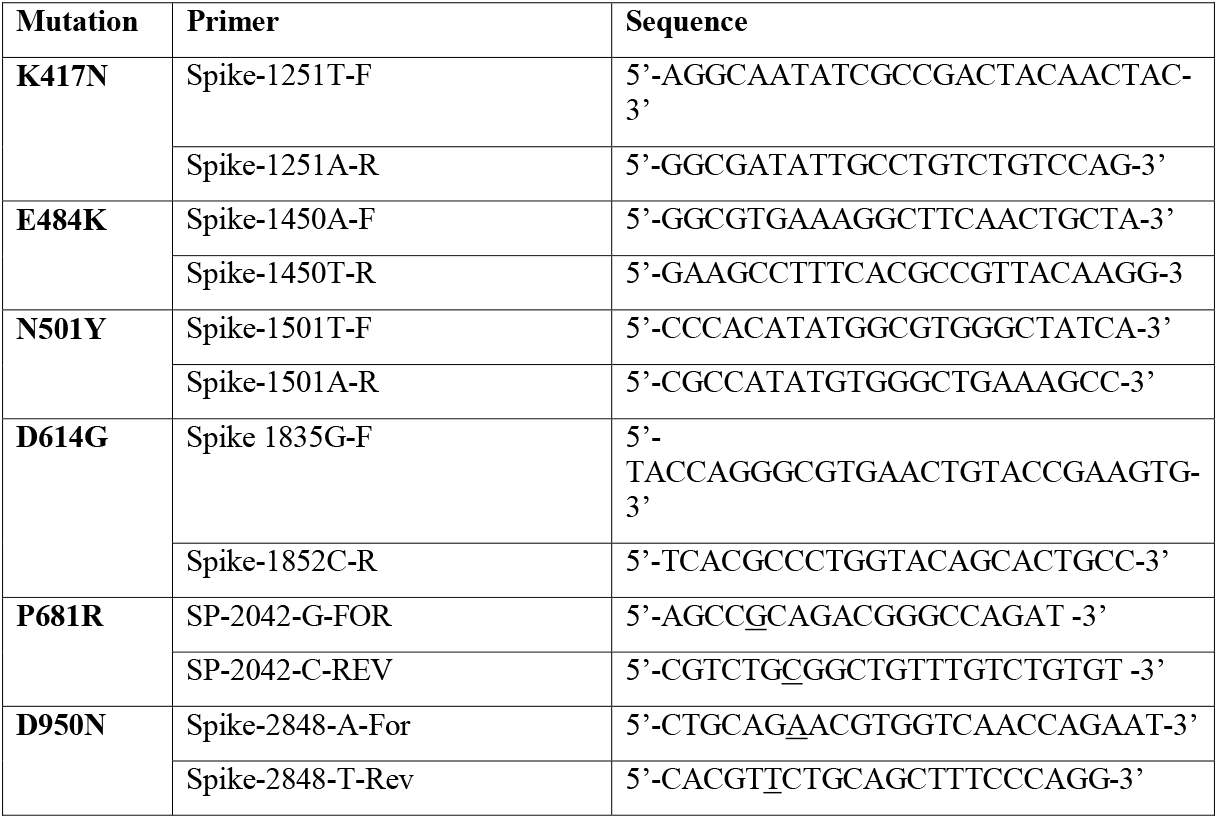
PCR primers for mutagenesis

To generate the Delta variant, we first introduced P681R and D950N mutations into the p278-1 spike expression clone with D614G. Next, to reconstruct the complete Delta variant spike, a synthesized DNA fragment (Genscript) corresponding to nucleotide positions 31-1446 and that includes mutations T19R, T95I, G142D, E156G Del157/158, L452R, and T478K of the Delta variant was spliced into the spike expression vector by HiFi DNA assembler cloning (NEB). The plasmid was prepared and verified as described above.

To express Omicron BA.1, a complete codon-optimized Omicron variant BA.1 spike was synthesized and inserted into pcDNA3.1 expression vector by Genscript. The clone includes the following amino acid differences from the Wuhan strain: A67V, Del69/70, T95I, G142D, Del143/145, N211I, insertion of EPE at 214, G339D, S371L, S373P, S375F, K417N, N440K, S477N, T478K, E484A, Q493R, G496S, Q498R, N501Y, Y505H, T547K, D614G, H655Y, N679K, P681H, N764K, D796Y, N856K, Q954H, N969K, and L981F. Similar to the spike variants based on the codon-optimized SARS-CoV-2 spike clone, p278-1, added at the carboxyl terminus is a 3xFlag-tag for identification of the spike protein by Western blot. This plasmid was also prepared and verified as described above.

Ten microliters of pseudovirus were incubated with serial dilutions of the serum samples for 1 hour at 37°C. Next, 100 μL of sera-pseudovirus were added to 293T-hACE2 cells in 96-well poly-D-lysine coated culture plates. Following 48 hours of incubation in a 5% CO2 environment at 37°C, the cells were lysed with 100 μL of Promega Glo Lysis buffer for 15 minutes at RT. Finally, 50 μL of the lysate were added to 50 μL luc substrate (Promega Luciferase Assay System). The amount of luciferase was quantified by luminescence (relative luminescence units (RLU)), using the Luminometer (Biosynergy H4). The percentage (%) virus inhibition was calculated as

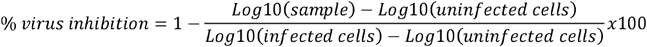

Serum from vaccinated mice was also compared by their 50% inhibitory dilution (IC50), defined as the serum dilution at which the virus infection was reduced by 50% compared with the negative control (virus +□cells).

### SARS-CoV-2

SARS-CoV-2 (USA-WA1/2020) was obtained from the University of Texas Medical Branch World Reference Center for Emerging Viruses and Arboviruses. SARS-CoV-2 virus Beta variant in lineage B. 1.351 was obtained from BEI Resources through the Africa Health Research Institute (NR-54009 BEI Resources, NIAID, NIH: SARS-Related Coronavirus 2, Isolate hCoV-19/South Africa/KRISP-K005325/2020, NR-54009, contributed by Alex Sigal and Tulio de Oliveira.). The virus was grown on Vero CCL81, aliquoted, and titrated by standard plaque assay on Vero E6 cells (Cercopithecus aethiops, Vero 76, clone E6, ATCC CRL-1586) as previously described (https://wwwnc.cdc.gov/eid/article/26/6/20-0516_article). The viral stock sequence was verified using Oxford Nanopore sequencing technologies using the NEBNext ARCTIC protocol.

### Plaque reduction neutralization test (PRNT)

Vero E6 cells were maintained in Dulbecco’s Modified Eagle Medium (DMEM) supplemented with 10% fetal bovine serum (FBS) and 100□U/mL penicillin-streptomycin. The assay was performed in duplicate using 12-well tissue culture plates in a biosafety level 3 facility. Serial dilutions of each serum sample were incubated with 55 plaque-forming units of SARS-CoV-2 or the Beta variant for 1□h at 37□°C. The virus-serum mixtures were added onto pre-formed Vero E6 cell monolayers and incubated for 1□h at 37□°C in a 5% CO2 incubator. The cell monolayer was then overlaid with 1% agarose:2X MEM:complete DMEM in a 1:1:1 ratio and incubated for 46 hours, after which time the plates were fixed and stained. Antibody titers were defined as the highest serum dilution that resulted in a >50% (IC50) reduction in the number of virus plaques. This method has previously been extensively validated on infected and control sera.

## Results

Preclinical studies were performed in eleven to thirteen-week-old female BALB/c mice, using yeast-produced SARS-CoV-2 RBD subunit vaccine candidates. Formulations were prepared with two protein amounts (2.3 and 7 μg). Similar to previously published experiments, we adsorbed the protein on aluminum hydroxide (alum) containing 100 μg aluminum per dose [17], and added, 20 μg CpG1826 just before injection. The animals were immunized twice intramuscularly on days 0 and 21. On day 35, the study was terminated, and serum and spleen samples were collected. Serum was tested for total anti-RBD IgG, and IgG subtypes, IgG2a and IgG1. Cytokines secreted into the supernatant upon restimulation of splenocytes were measured. Sera were further evaluated for neutralizing antibodies against the original SARS-CoV-2 and the variants-of-concern.

### RBD219-N1C1/alum + CpG triggers a robust humoral immune response in mice

Mice that received the vaccine antigen formulated with alum alone showed above baseline IgG titers in all groups (**Figure 1A**). The magnitude of the response was dose-dependent, as evidenced by the observation that mice that had received 7 μg of RBD antigen showed mean IgG titers approximately 100x higher than mice that only received 2.3 μg of the RBD antigen. The addition of 20 μg CpG to the vaccine formulation significantly raised geometric mean titers by an additional factor of approximately 100 (when comparing formulations with a constant amount of 7 μg of RBD antigen).

**Figure 1:**
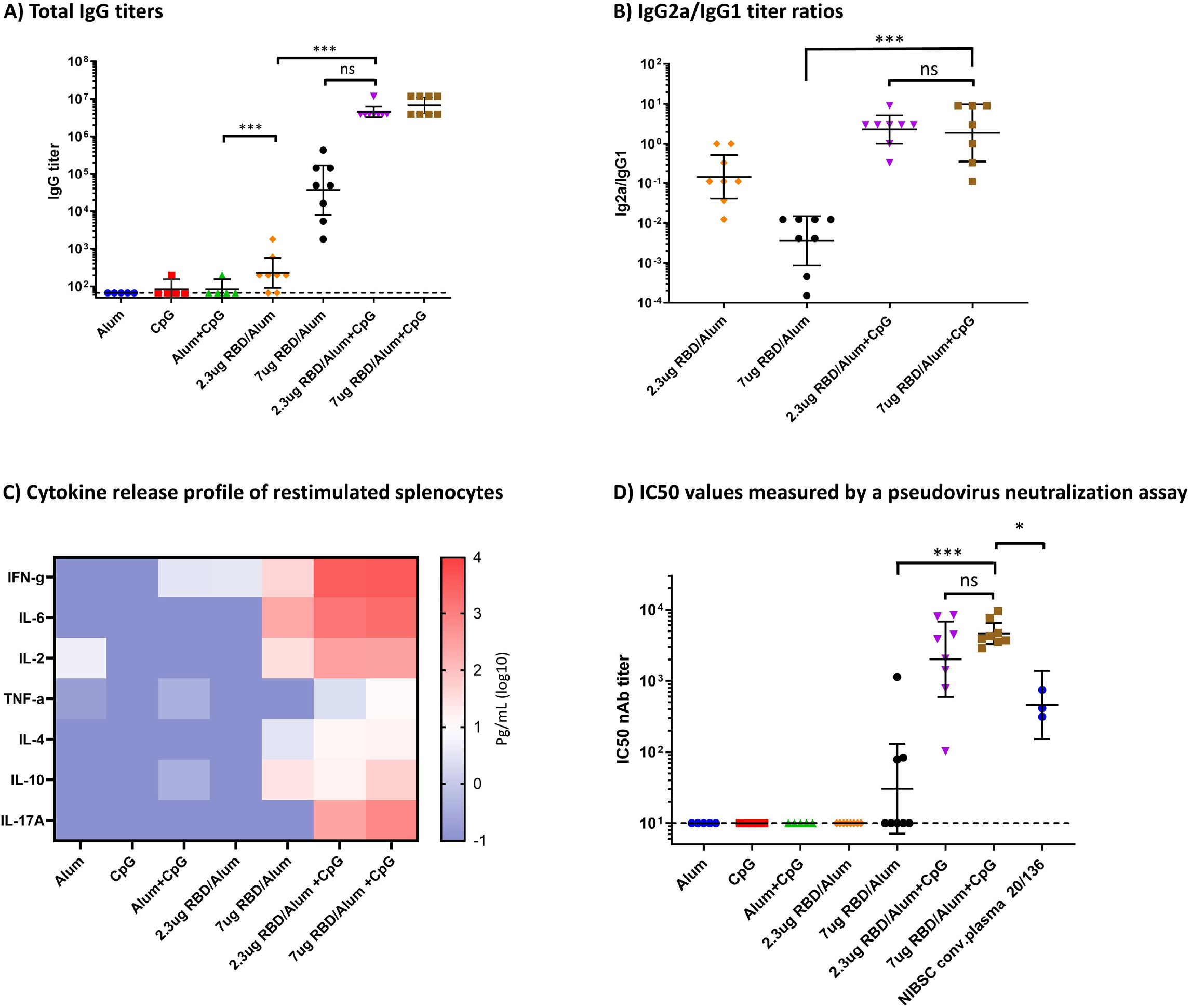
Humoral and cellular immune response in immunized mice. **A)** Total IgG titers at day 35, measured by ELISA. **B)** IgG2a/IgG1 titer ratios. Geometric mean and 95% confidence intervals are shown. **C)** Heatmap of secreted cytokine levels measured in the supernatants from splenocytes re-stimulated with the SARS-CoV-2 RBD protein. Median values were calculated within each treatment group for each cytokine and plotted. **D)** IC50 values measured by a neutralization assay using a lentiviral SARS-CoV-2 Wuhan pseudovirus. Mann-Whitney tests were performed to evaluate for statistical significance between different groups. p> 0.12(ns), p < 0.033 (*), p<0.001 (***).

We did not observe any statistically significant differences between the mean IgG titers in the CpG-immunized groups based on the amount of antigen delivered. As expected for vaccines adjuvanted with alum alone, IgG1 titers were greater than IgG2a titers (**Figure 1B**). The addition of CpG compensated for the alum-induced Th2-bias, producing a balanced IgG2a to IgG1 antibody response.

### RBD219-N1C1/alum + CpG elicits a strong cellular immune response in mice

Cytokines secreted into the supernatant from splenocytes restimulated using recombinant SARS-CoV-2 RBD219-N1protein were analyzed in a Luminex assay (**Figure 1C**). As expected for vaccines adjuvanted with alum alone, a Th2 cytokine response was induced, while the addition of CpG to the formulation raised a Th1/Th2 balanced cytokine response, with increased levels of anti-inflammatory cytokines IL-4 and IL-10, antiviral interferon IFN-g, and proinflammatory cytokines IL-6 and IL-2. We also observed that for the RBD/alum formulation the cytokine secretion increased significantly when the protein doses increased from 2.3 to 7 μg, whereas for the RBD/alum + CpG formulations the dose-dependent release of cytokines was much less prominent.

### RBD219-N1C1/alum + CpG elicits a strong neutralizing antibody response in mice against the original Wuhan SARS-CoV-2

Sera were tested for neutralizing antibodies in a pseudovirus assay. **Figure 1D** shows the IC50 values measured by a neutralization assay using a lentiviral SARS-CoV-2 pseudovirus with a spike protein closely mimicking the sequence of the spike protein of the first virus isolated in Wuhan, China (Genbank entry MN908947.3). We detected no neutralizing antibodies in the sera of mice immunized twice with 2.3 μg RBD/alum, and only 3 out of 8 mice immunized with two doses of the 7 μg RBD/alum showed neutralizing titers. In the latter group, we also observed a more widespread response difference between the individual mice.

The addition of 20 μg CpG to the vaccine formulation raised the mean neutralizing titers by a factor of approximately 100. Notably, sera from mice that received 7 μg RBD/alum+CpG showed on average 5.5 times higher neutralizing antibody titers compared to the NIBSC 20/136 human convalescent plasma standard. There was no statistical difference between the amount of neutralizing antibodies when the RBD219-N1C1 protein was increased from 2.3 to 7 μg, however, we did observe a reduction in the geometric mean titers (GMT) and the intra-cohort variation (Comparing 7 μg RBD/Alum+CpG GMT=4631 95% confidence interval 3293-6514, to 2.3 μg RBD/Alum+CpG GMT=2012, 95% confidence interval 595-6808).

### A booster dose may not require the addition of CpG to elicit strong humoral and cellular immune responses in mice

We evaluated the need for CpG to be included in the prime/boost vaccination strategy. As can be seen in **Figures 2A-B**, when the second dose of RBD219-N1C1/alum+CpG was replaced with a formulation containing RBD219-N1C1/alum alone (same formulation without CpG), no significant drop in the antibody titers were observed when compared to two doses of RBD219-N1C1/alum+CpG. We also detected similar amounts of neutralizing antibodies (**Figure 2D**), indicating that a boost without CpG may be sufficient to induce a protective immunological response. The presence or the absence of CpG in the second dose of the vaccine did however impact the cytokine profile (**Figure 2C**). The median values of secreted INF-g, IL-6 and IL-17A were increased by the addition of CpG to the booster, while the median value of TNF-a decreased. In conclusion, the neutralizing antibody titers are not impacted by absence of CpG in the booster dose, but the cellular profile starts skewing slightly more towards a Th2 type response.

**Figure 2:**
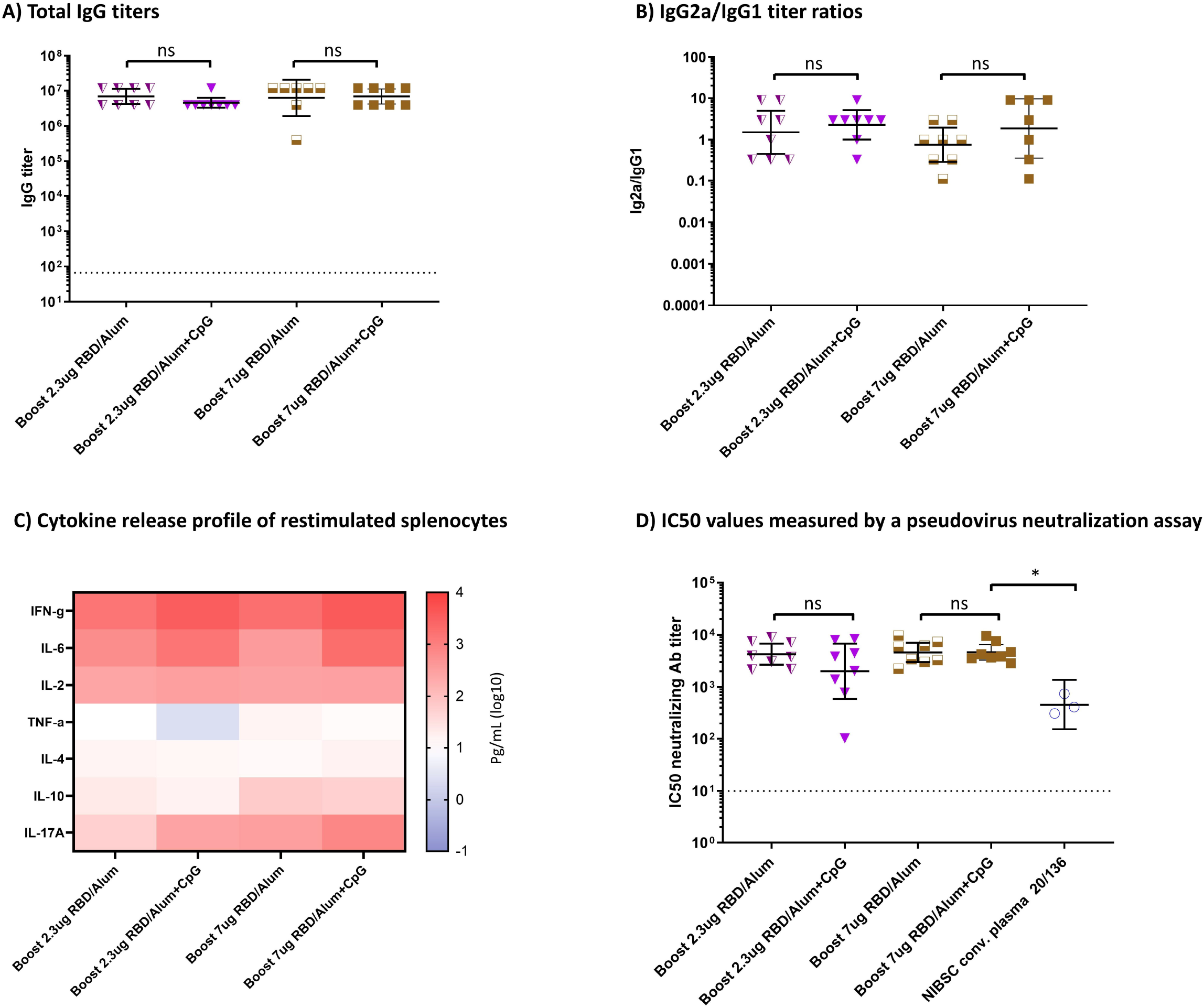
Comparison of RBD219-N1C1-specific immune responses after one dose with CpG and a boost without CpG versus two vaccine doses with CpG. Geometric mean and 95% confidence intervals are shown. **A)** Total IgG titers measured from mouse sera against RBD protein, **B)** IgG2a/IgG1 titer ratios from mouse sera against RBD protein, **C)** Heatmap of secreted cytokines measured in supernatants from splenocytes re-stimulated with RBD protein. **D)** IC50 values measured by a neutralization assay using a lentiviral Wuhan SARS-CoV-2 pseudovirus. Mann-Whitney tests were performed to evaluate for statistical significance between different groups. p> 0.12(ns), p < 0.033 (*). The dotted lines indicate the limit of quantification.

### The RBD219-N1C1/alum+CpG vaccine elicits robust neutralizing antibodies effective against the B.1.1.7 (Alpha) and B.1.351 (Beta) SARS-CoV-2 variants

To determine if the vaccine would cross-protect against different variants of concern and a variant of interest of the SARS-CoV-2 virus, we evaluated the neutralizing antibodies from selected pooled sera against four pseudoviruses: (i) Original Wuhan SARS-CoV-2 isolate, (ii) B.1.1.7 (Alpha) SARS-CoV-2 variant, (iii) B.1.351 (Beta) SARS-CoV-2 variant (**Figure 3A, Supplementary Figure 1, Supplementary Table 1**).

**Figure 3:**
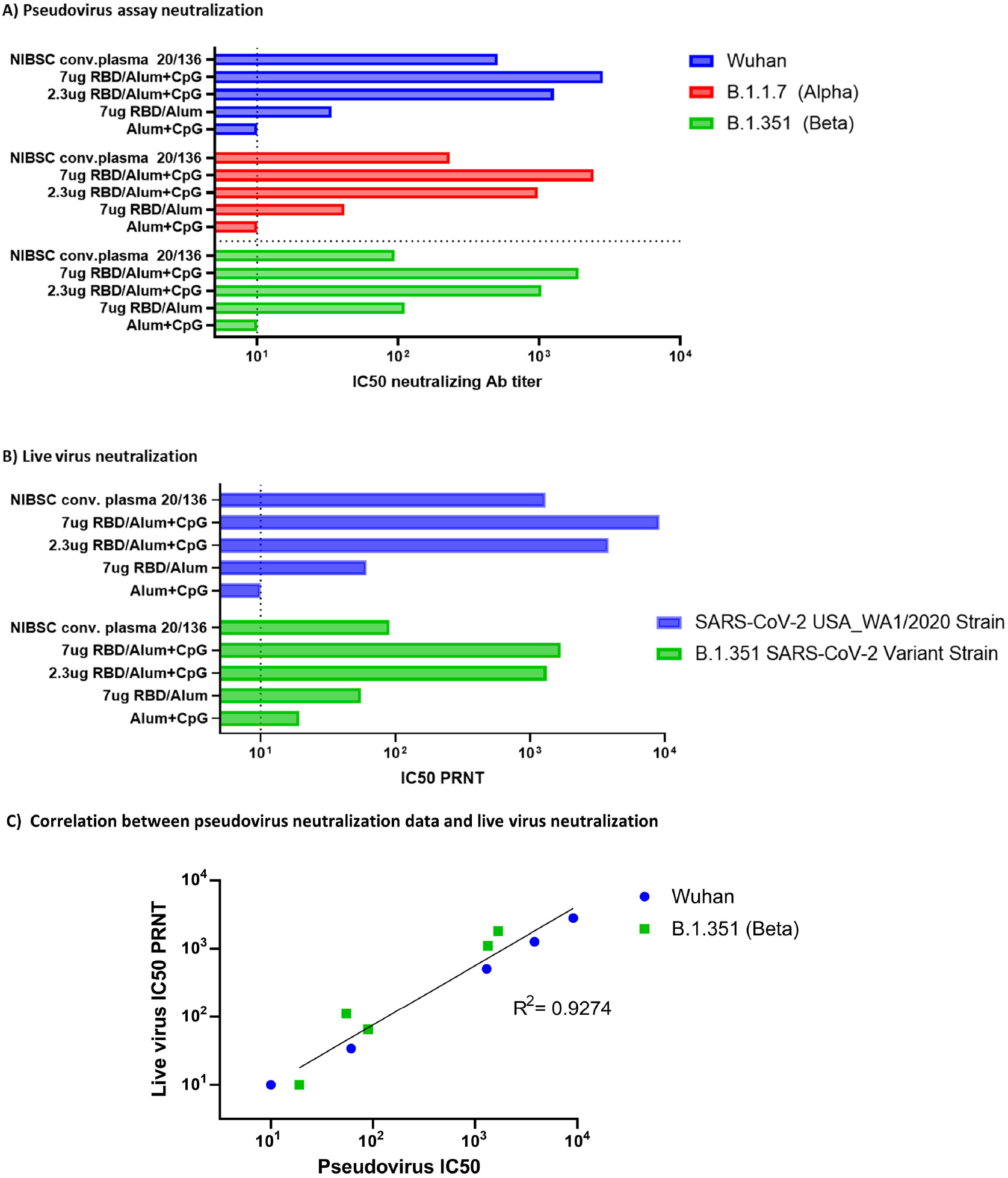
Neutralizing activity by sera from vaccinated mice against Alpha and Beta variants of the SARS-COV-2 pseudovirus. **A)** Neutralization of various pseudovirus mimicking different variants of concern: Wuhan SARS-CoV-2 (blue), B.1.1.7 (Alpha) SARS-CoV-2 variant (red), B.1.351 (Beta) SARS-CoV-2 variant (green). The bars graph represents the geometric means of technical replicates (n=6, except for the group vaccinated with 7ug RBD/Alum n=2). The dotted line indicates the limit of quantification. **B)** Live virus neutralization (PRNT assay): Original SARS-CoV-2 (blue), B.1.351 (Beta) SARS-CoV-2 variant (green). The dotted line indicates the limit of quantification. The bars graph represents the geometric means of technical replicates (n=2). **C)** Correlation graph, comparing IC50s of the live virus and pseudovirus neutralization.

Sera from mice vaccinated with the CpG adjuvanted vaccine candidate maintained the ability to neutralize pseudoviruses which include key mutations, mimicking important variants of interest and concern. Comparing the average IC50 values, the 7 μg RBD219-N1C1/alum+CpG vaccine showed only 1.6 times lower values against the B. 1.351 (Beta) pseudovirus compared to the Wuhan, pseudovirus (**Figure 3A**). Meanwhile, in the same assay, the Human convalescent plasma (WHO International Standard for anti-SARS-CoV-2 immunoglobulin, NIBSC code: 20/136 [32]) showed a 7.8 fold reduction in its neutralization ability of the Beta pseudovirus.

To verify the results obtained by the pseudovirus experiments, we also evaluated the virus-neutralizing ability of the pooled mouse sera, using isolated live viruses in a plaque reduction neutralization assay (PRNT) **(Figure 3B, Supplementary Figure 2, Supplementary Table 2)**. The results of the live virus assay were concordant with the results found in the pseudovirus assay **(Figure 3C)**. For example, against the B.1.351 live virus, the IC50 values for the mouse serum obtained after two vaccinations with 7 μg RBD219-N1C1/alum+CpG are 18-fold higher than the human convalescent plasma standard (**Figure 3B**). This increased protection against B.1.351 is similar to the 20-fold increased protection calculated from the pseudovirus data. The equivalent results of the two isolated live strains of SARS-CoV-2 demonstrate that our pseudovirus-based neutralization assay can be used as a valid alternative to more rapidly screen for neutralizing activity of the serum antibodies against the spike protein of various SARS-CoV2 variants of interest. The data reported here reinforces that the RBD219-N1C1/alum+CpG vaccine formulation is potentially suitable for broad neutralization against SARS-CoV-2 including variants-of-concern.

### The RBD203-N1/alum+CpG vaccine elicits robust neutralizing antibodies effective against the B.1.617.2 (Delta) and B.1.1.529 (Omicron) SARS-CoV-2 variants

Mice immunized with the RBD203-N1/alum +CpG vaccine produced similarly efficacious neutralizing antibodies against the Wuhan pseudovirus as mice vaccinated with RBD219-N1C1 formulations (IC50 values of 2699 and 2827, respectively) **(Figure 4, Supplementary Figure 3, Supplementary Table 3)**. The neutralizing titers dropped about 6.8-fold against the B.1.617.2 (Delta) variant (IC50: 392) and 57.2-fold against the B.1.1529 (Omicron) variant (IC50: 44).

**Figure 4:**
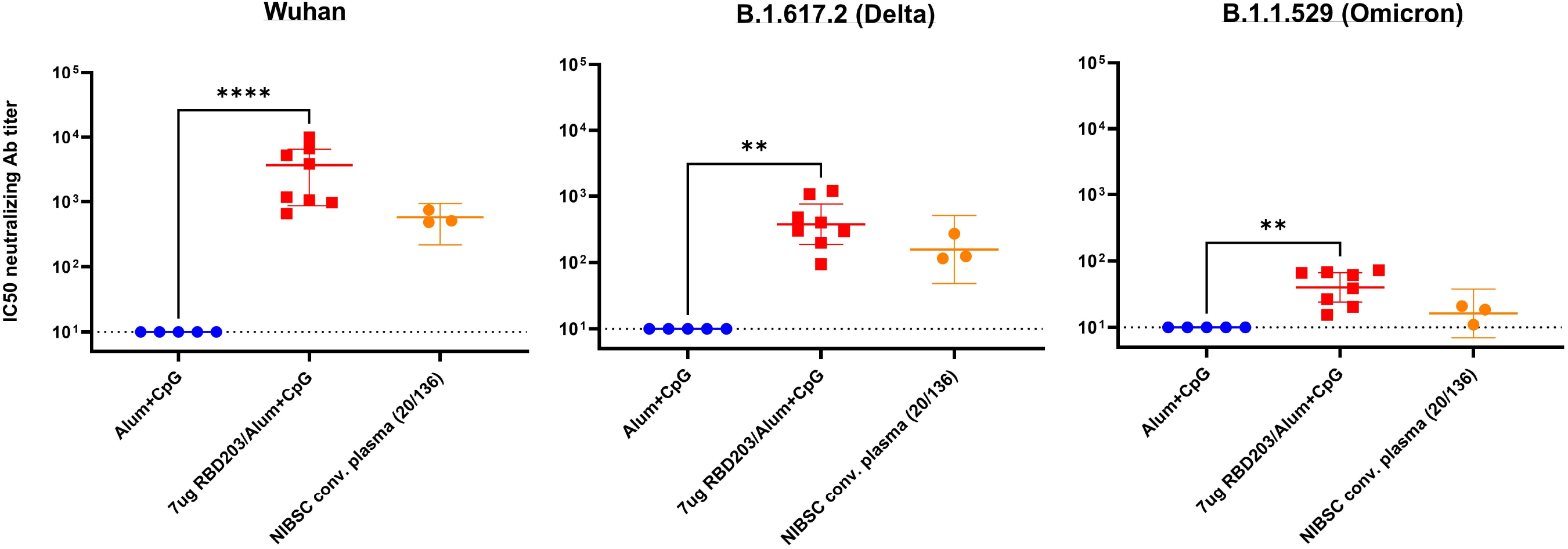
Neutralizing activity by sera from vaccinated mice against Delta and Omicron variants of the SARS-CoV-2 pseudovirus. Neutralization of various pseudovirus mimicking different variants of concern: Wuhan SARS-CoV-2 (left panel), B.1.617.2 (Delta) SARS-CoV-2 variant (middle panel), B. 1.351 (Omicron) SARS-CoV-2 variant (right panel). The human NIBSC convalescent plasma was included as a positive control, serum from mice vaccinated with alum+CpG was included as a negative control. Data points are averages of technical replicates (n=2). The dotted line indicates the limit of quantification p < 0.0021 (**), p<0.0001 (****).

## Discussion

Here we show the results of preclinical studies in mice immunized with SARS-CoV-2 recombinant RBD protein vaccine candidates produced in yeast and formulated with aluminum hydroxide (alum) and CpG deoxynucleotides, a formulation comparable to the formulation of the Corbevax vaccine that recently received emergency use authorization by Indian Regulators.

Studies have shown that neutralizing antibody titers are a good correlate of protection against SARS-CoV-2. While we recognize the limitations of directly comparing human and mouse sera samples, we here used convalescent human serum as a benchmark of natural immunization through infection. The RBD antigen, adjuvanted with alum and CpG, elicited neutralization antibody titers against the Wuhan SARS-CoV-2 in a pseudovirus assay that were significantly higher than those in sera from mice vaccinated with RBD/alum alone and also higher than in the NIBSC 20/136 human convalescent plasma standard. In addition, the RBD219-N1C1/alum+CpG vaccine provided superior protective antibodies against emerging variants-of-concern. In our assay, human convalescent plasma showed a 7.8-fold reduction of the neutralization of the B.1.351 pseudovirus (IC50=65) compared to the Wuhan pseudovirus (IC50=508). By contrast, sera from mice vaccinated with 7 μg RBD219-N1C1/alum/CpG only showed a 1.6-fold reduction (IC50=1811 and IC50=2827), potentially indicating broader protection offered by the vaccine compared to natural infection. Furthermore, when comparing our neutralizing data to the results of (pre)clinical studies from other SARS-CoV-2 vaccines, we observe that our vaccine candidate induces among the highest titers of neutralizing antibody titers [33, 34].

While we observed an expected drop of the neutralizing antibodies against the Delta (6.8 fold) and Omicron variant (57.2 fold), the mouse sera elicited by vaccination with the ancestral Wuhan RBD did provide significant antibodies to protect against possible mutant infections. For comparison, for the ALFQ-adjuvanted Spike-nanoparticle, a 79.3 fold reduction of the ID50 against omicron-pseudoviru was reported with non-human primate serum [35].

This level of protection against variants-of-concern was also seen with live viruses. Against B.1.351 live virus, the IC50 neutralization values of mouse serum obtained after two vaccinations with 7 μg RBD219-N1C1/alum+CpG (IC50PRNT=1678) are 18-fold higher than the human convalescent plasma standard (IC50PRNT= 90). This again emphasizes the broader protection offered through the antibodies produced by a recombinant protein-based vaccine versus the antibodies induced by a natural infection.

Multiple vaccines containing CpG adjuvants [36] have undergone evaluation in human clinical trials, including vaccines for cancer, allergy, and infectious diseases. A recent phase 1 study of the SARS-CoV-2 trimeric spike protein with alum+CpG showed excellent tolerability and immunogenicity [37]. While our RBD219-N1C1 antigen formulated only with alum elicited neutralizing antibody titers against SARS-CoV-2 that exceeded titers of the human convalescent plasma standard [17], there was a higher level of variability, together with a stronger bias towards Th2-type immunity.

We further evaluated the possibility of obtaining high neutralizing antibody titers with a single dose of the CpG adjuvanted vaccine, followed by a boost with RBD/alum. While the addition of CpG to the formulation of the prime vaccine resulted in an approximately 100-fold increase in antibody titers, including CpG in the formulation of the boost vaccine induced only a minimal additional immunological effect. The addition of CpG did have a limited Th1-skewing effect based on the cytokine release profile of restimulated splenocytes. Therefore, future formulations of the booster vaccines may still require CpG, but in much lower amounts, thus reducing supply chain stress and reducing cost.

Further studies are in progress to examine whether the cross-protection of the vaccine corresponds to the enhanced neutralizing breadth phenomenon noted with previous COVID-19 infection followed by mRNA vaccine boosters [38]. Additionally, an important shortcoming of this initial investigation is that we did not address the duration of protection achieved after boosting with RBD/alum alone. While antibody titers and cytokine profiles were in line with those observed in mice that received two vaccinations, we do not know if that will also hold after an extended period Additional long-term studies are underway to evaluate the duration of the response. In parallel studies, we are also expanding our evaluation of a one-dose approach, which is of great interest to the scientific community since it would improve the ease of delivery as well as confidence and compliance with the necessary vaccination protocol.

Even though there have been several vaccines developed against COVID-19, there remains an urgent need to provide additional tools against a virus that continues to spread, specifically in LMICs. There are several requirements that a COVID-19 vaccine for impoverished populations must meet. The vaccine must be able to be produced on a large scale. Moreover, storage and delivery logistics of the vaccine must be compatible with the infrastructure in LMICs (i.e., without the need for ultralow temperature storage). The yeast-produced RBD219-N1C1 vaccine uses a production platform very similar to that used for the hepatitis B vaccine and fulfills these criteria. It can be produced at a low cost, with high yields, and by manufacturers that have previous experience with yeast-based recombinant protein vaccines.

## Acknowledgements

We are grateful to Dr. Vincent Munster (NIAID) for providing the spike expression plasmid for SARS-CoV-2. We like to thank Deborah Higgins, Dr. Fred Cassels, and Dr. Bob Sitrin (PATH) for their valuable feedback on our data and the manuscript. This work was supported by the following funding sources: Robert J. Kleberg Jr. and Helen C. Kleberg Foundation, USA; Fifth Generation, Inc. (Tito’s Handmade Vodka), USA; JPB Foundation, USA, National Institute of Health-National Institute of Allergy and Infectious Diseases (NIH/NIAID), USA, Grant number AI14087201); Baylor College of Medicine; and Texas Children’s Hospital Center for Vaccine Development Intramural Funds, USA.

## Author Contributions

JP contributed to the design and planning of the experiments, prepared the vaccine formulations, quantified neutralizing antibodies, processed all data, prepared the figures, and drafted the manuscript.

US contributed to the design and planning of the experiments, the evaluation, and the interpretation of the data. US also drafted parts of the original manuscript and edited subsequent versions of the manuscript.

BZ and JW established the expression of recombinant protein RBD-219-N1C1 in yeast. ZL and

JL performed the fermentation and purification of recombinant protein RBD219-N1C1.

WHC, RTK, and RA contributed to the characterization of the recombinant protein RBD219-N1C1.

LV contributed to the design and the planning of the experiments, the execution and data analysis of ELISA and Luminex experiments, and the review and editing of the manuscript.

BK contributed to the design and planning of the experiments and performed all animal manipulations.

PG contributed to the design and planning of the experiments, the evaluation, and the interpretation of the data.

BL and JTK prepared the plasmids for the pseudovirus-based neutralization assays.

MJV, SRT, and CP have executed the pseudovirus experiments and processed data.

SR executed the live SARS-CoV-2 virus assays inside the BSL-3

MR and VP contributed to the design and planning of the experiments and managed the production of RBD219-N1C1 at Biological E, Limited (Hyderabad, India).

MEB and PJH conceptualized and designed the experiments and managed the process of manuscript review and editing.

All authors contributed equally and provided critical feedback, reference sources, and critical revisions for intellectual content, and verified the information presented here.

## Supplementary Information

Supplementary Information was added as a separate document.

## Data Availability Statement

The data that support the findings of this study are available on request from the corresponding author.

