## Supplementary for "Receptor-binding domain recombinant protein on alum-CpG induces broad protection against SARS-CoV-2 variants of concern"

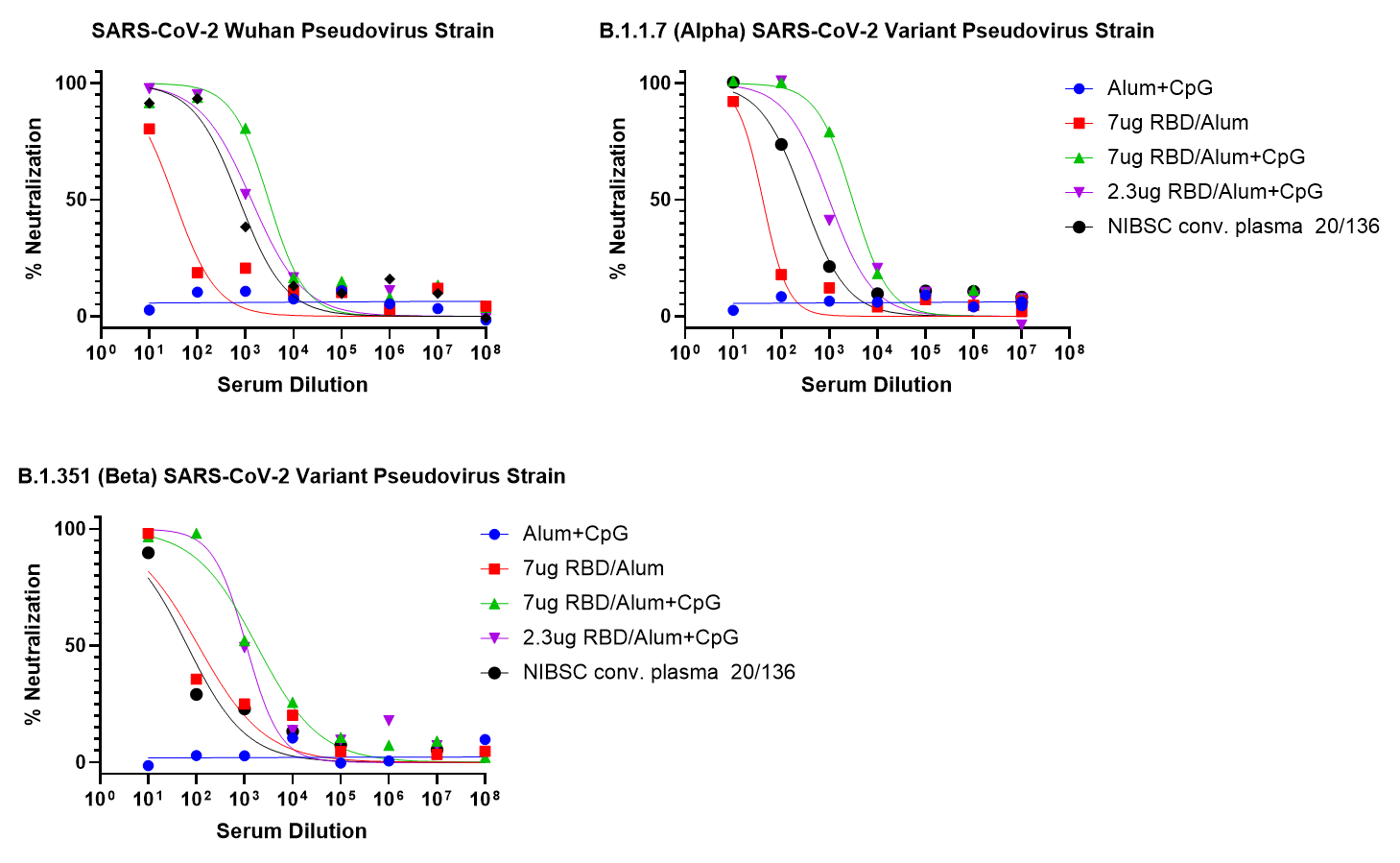

**Supplementary Figure 1 : Neutralization of SARS-CoV-2 Pseudovirus mimicking different variants of concern. Data points are averages of technical replicates (n=3)**

**Supplementary Table 1: Mean IC50 values of pseudovirus neutralization assays**

|  | **Pseudovirus IC50** | | |
| --- | --- | --- | --- |
|  | **Wuhan** | **N501Y** | **K417N-E484K-N501Y** |
| NIBSC Conv plasma | 508 | 294 | 65 |
| 7ug RBD/Alum+CpG | 2827 | 3060 | 1811 |
| 2.3ug RBD /Alum+CpG | 1267 | 994 | 1101 |
| 7ug RBD/Alum | 34 | 41 | 111 |
| Alum+CpG | 10 | 10 | 10 |

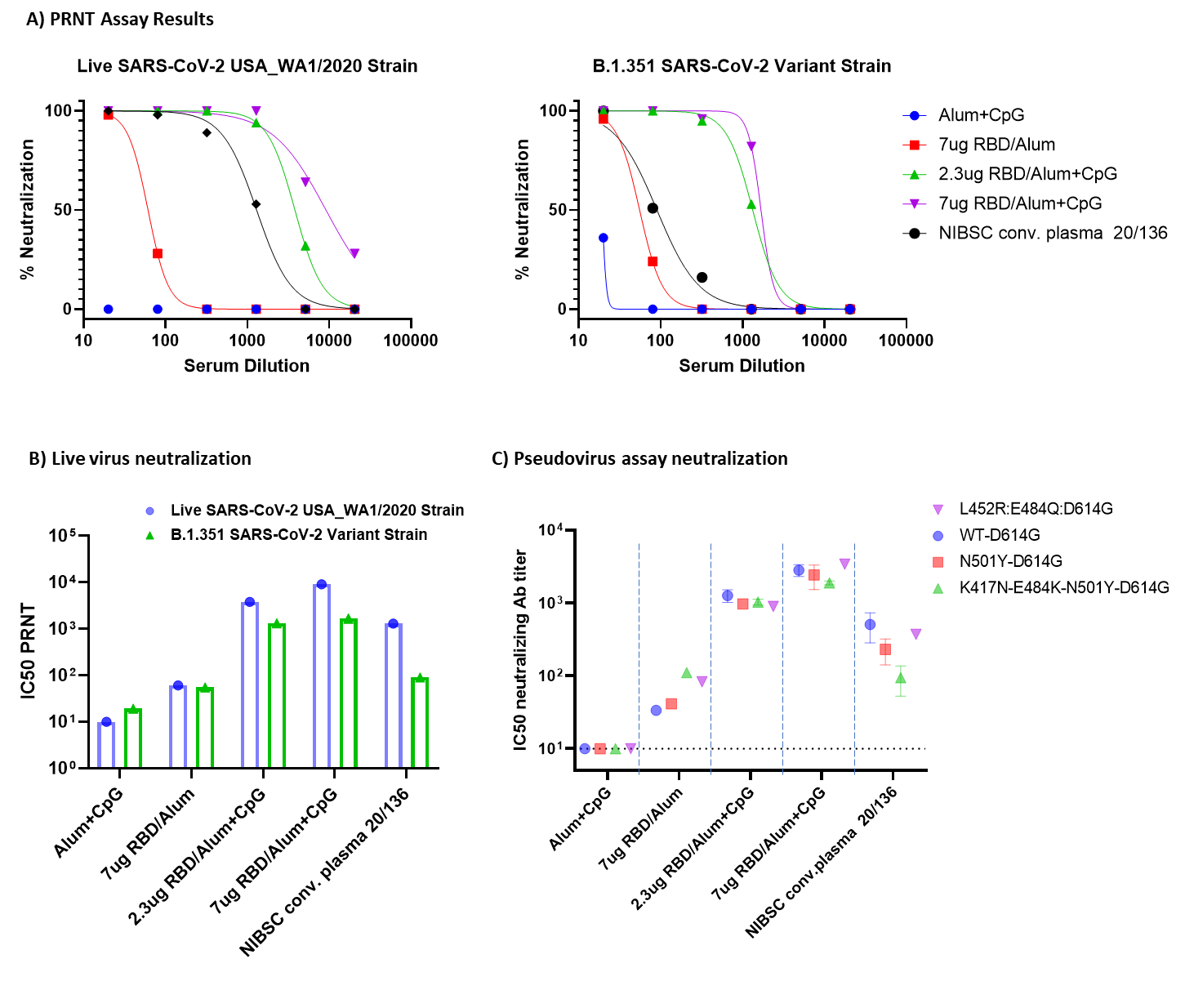

**Supplementary Figure *2*: Live Virus N****eutralization.** A) Plaque reduction neutralization test (PRNT) with select serum samples using the live virus strain USA_WA1/2020, isolated before the emergence of key mutations, and PRNT with select serum samples using a live virus strain of the South African variant (B.1.351). Datapoints are averages of technical replicates (n=2)

**Supplementary Table 2: Mean IC50 values of pseudovirus neutralization assays**

|  | **Live virus IC50 PRNT** | |
| --- | --- | --- |
|  | **Wuhan** | **B.1.351** |
| NIBSC Conv plasma | 1296 | 90 |
| 7ug RBD/Alum+CpG | 9130 | 1678 |
| 2.3ug RBD /Alum+CpG | 3810 | 1328 |
| 7ug RBD/Alum | 61 | 55 |
| Alum+CpG | 10 | 19 |

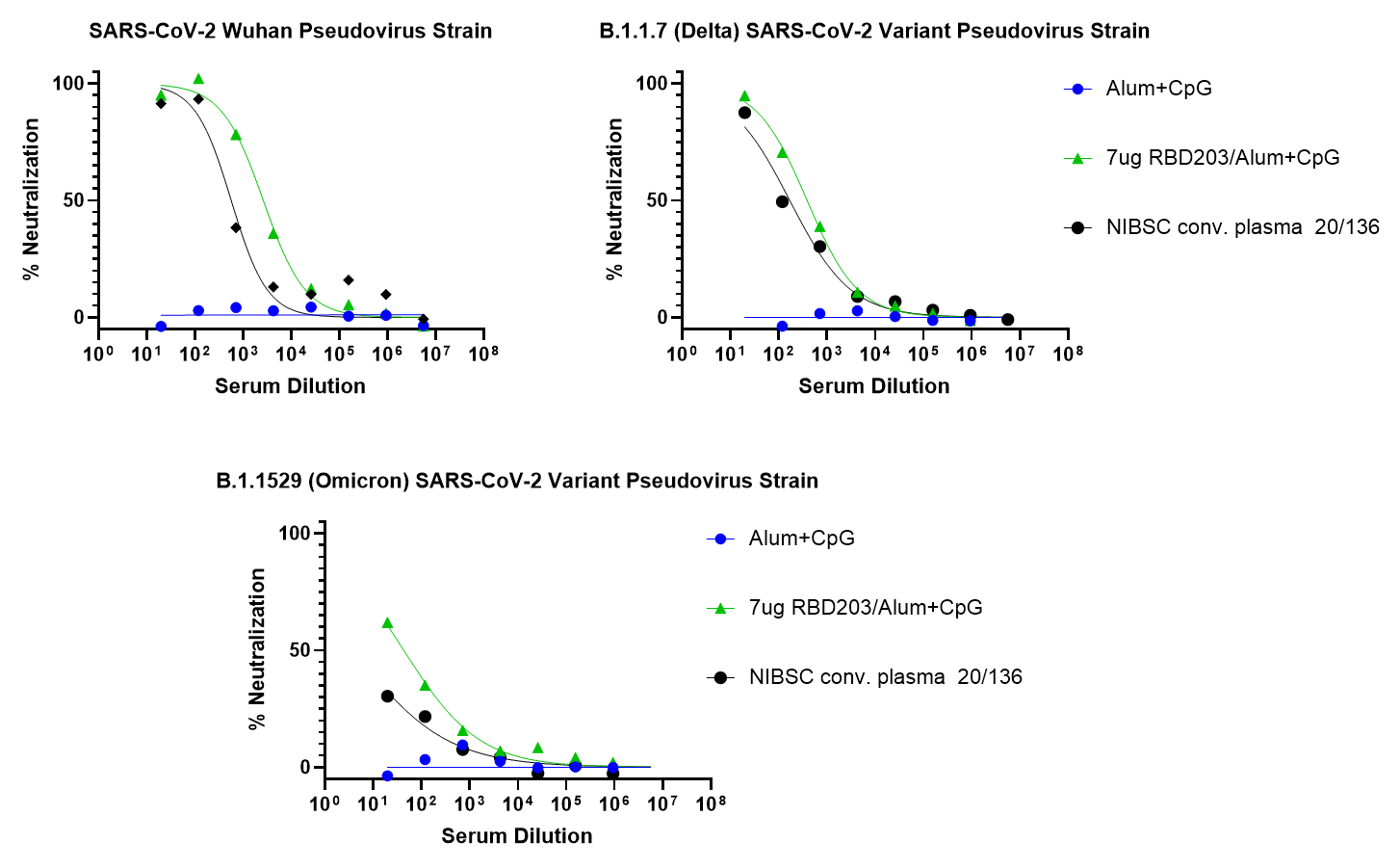

**Supplementary Figure 3: Neutralization of SARS-CoV-2 Pseudovirus mimicking different variants of concern. Data points are averages of technical replicates (n=2) of individual mice (n=8).**

**Supplementary Table 3: Mean IC50 values of pseudovirus neutralization assays for RBD203-N1**

|  | **Pseudovirus IC50** | | |
| --- | --- | --- | --- |
|  | **Wuhan** | **B.1.617.2 (Delta)** | **B.1.1529 (Omicron)** |
| NIBSC Conv plasma | 572 | 171 | 10 |
| 7ug RBD203/Alum+CpG | 2699 | 392 | 44 |
| Alum+CpG | 10 | 10 | 10 |
